# On the completeness of existing RNA fragment structures

**DOI:** 10.1101/2024.05.06.592843

**Authors:** Xu Hong, Jian Zhan, Yaoqi Zhou

**Affiliations:** Institute of Systems and Physical Biology, Shenzhen Bay Laboratory, Shenzhen 518055, China; Ribopeutic Inc, Guangzhou International Bio Island, Guangdong, 510320, China

**Keywords:** RNA fragment, RNA modeling, RNA structure, RNA pseudo-torsion angle

## Abstract

Success in protein structure prediction by the deep learning method AlphaFold 2 naturally gives arise the question if we can do the same for RNA structure prediction. One reason for the success in protein structure prediction is that the structural space of proteins at the fragment level has been nearly complete for many years. Here, we examined the completeness of RNA fragment structural space at dimeric, trimeric, tetrameric, and pentameric levels. We showed that the RNA structural space is not even complete at the di-nucleotide level, whereas the exponential increase of new structural fragments is observed at tetrameric and pentameric levels. Moreover, the number of backbone fragments found in RNA (2510) is far smaller than the number of backbone fragments found in proteins (6652). This further suggests that a structural space currently observed in RNA is far from complete, considering that the RNA backbone (6 torsion angles) has more degrees of freedom than the protein backbone (3 torsion angles with one nearly fixed). In addition, we found that the three-atom representation (one backbone atom C4’ and two sidechain atoms C1’ and N1) has the lowest number of structural fragments, suggesting it as the most “stable” structural frame for building up the entire RNA structure.

## Introduction

The recent success of AlphaFold2 ^1^ for protein structure prediction relied on the large database of protein structures (∼200,000) in the protein databank ^2^. This dataset has long ago achieved nearly complete structure space at the fragment [3, 4-residue ^3^, 7, 9-residue ^4^ and even structural domain levels ^5^]. This nearly complete structural space allows the dominance of fragment-based assembly for de novo protein structure prediction prior to 2018 and paves the way for the success of end-to-end deep learning techniques such as AlphaFold 2 for high-accuracy prediction ^6^.

Like proteins, many RNAs have unique three-dimensional structures that are wholly determined by their sequences. These structured RNAs, including tRNA, rRNA, ribozymes, riboswitches, and long noncoding RNAs, are the keys for understanding their functional mechanisms ^7^. However, unlike proteins, RNAs are more challenging to be determined by nuclear magnetic resonance spectroscopy, X-ray crystallography, or cryogenic electron microscopy due to their unique physio-chemical properties ^7^. Currently, RNA-containing structures represent only 3% of all structures deposited in the protein databank ^2^. This raises the question if the current fragment-structural library contained in the protein databank is sufficient for AI-based learning ^8, 9^ or fragment-based assembly for RNA structure prediction. Fragment-based techniques have been employed for prediction of RNA 3D structures [e.g. Rosetta, SWM, FARFAR ^10^/ FARFAR2 ^11^, 3dRNA ^12, 13, 14, 15^, FebRNA ^16^], RNA-RNA complex structures ^17^, and protein-RNA complex structures ^18^.

Previously, RNA fragment structures were classified according to two pseudo-torsion angles θ (P_i_, C4’_i_, P_i+1_, C4’_i+1_) and η (C4’_i-1_, P_i_, C4’_i_, P_i+1_) ^19^. However, the conformations of two fragments can be significantly different even if they are in the same pseudo-torsion bin (e.g. a typically employed 5 degree bin). Thus, the use of two pseudo-torsion angles will likely underestimate the structural space of RNAs.

The completeness of a fragment structure library for proteins was measured according to the structural similarity between fragment backbone atoms, because protein structures can be characterized by regular secondary structure elements (helix and sheet) of backbones with sidechains as rotamers around main chains ^20^. RNA structures, on the other hand, are driven by base pairing of nucleotide bases whereas the backbone was considered as rotamers around bases ^21, 22^. Thus, the structural characterization of RNA fragments should consider both backbone and base atoms. Here, we investigated several reduced representations (backbone only, base only and mixed, as shown in Fig. 1) and found that the structural space of RNA fragments is not yet complete even at the di-nucleotide level.

**Figure 1.**
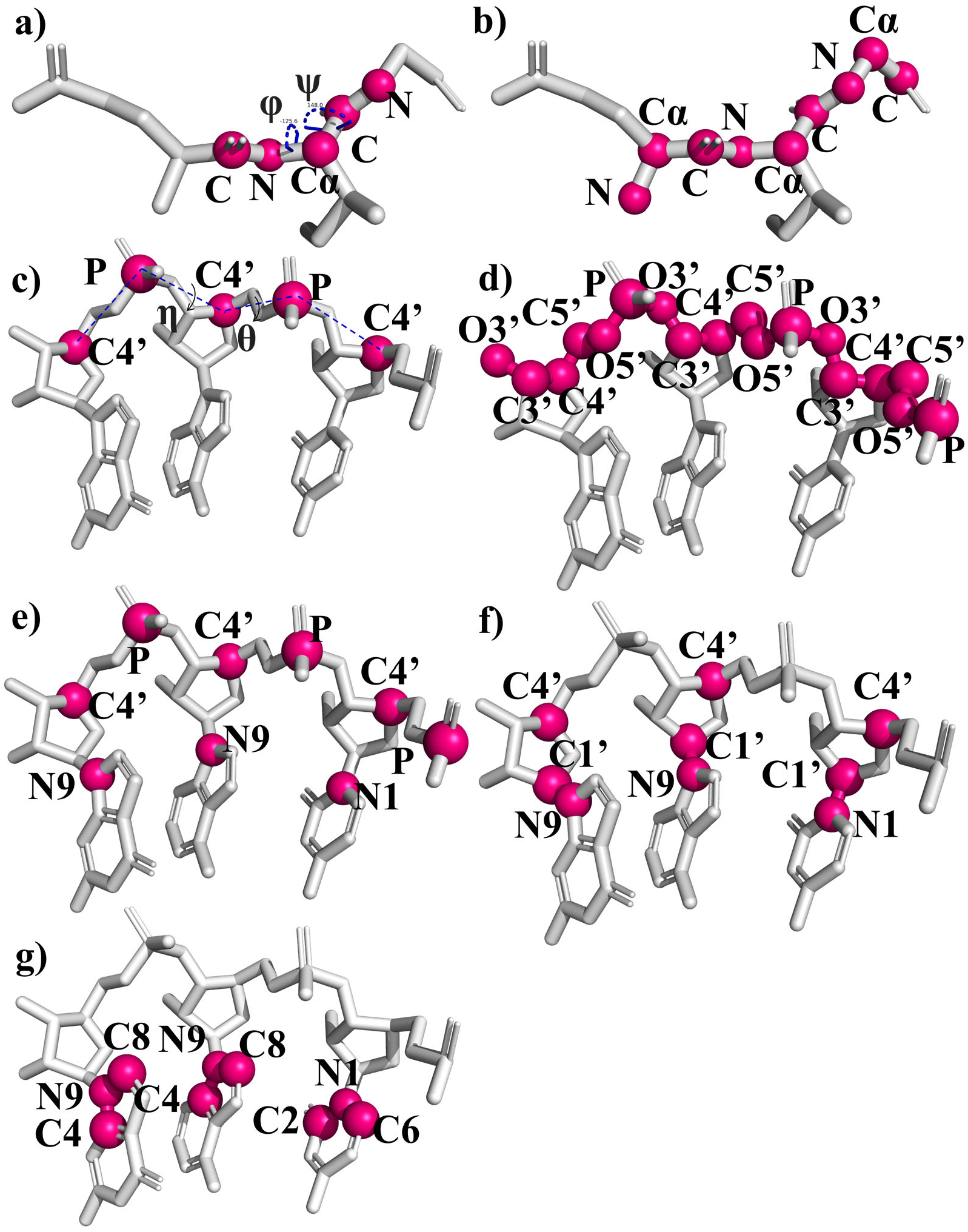
The atoms and torsion angles employed for protein and RNA fragment-structure classifications. a) Protein backbone torsion angles (ϕ, ψ). b) Protein: three mainchain atoms (N, Cα, C) of an amino acid residue commonly employed as a reference frame. c) Two pseudo-torsion angles (θ, η) for RNA backbone-structure characterization are rotated about the P_i_-C4’_i_ and C4’_i_-P_i+1_ pseudo-bonds, respectively. d-g) Different atoms from the backbone and/or the sidechains of an RNA used as a reference frame. They are B6S0 (d), B2S1 (e), B1S2 (f) and B0S3 (g) denoted according to the number of atoms from **B**ackbone or **S**idechain.

## Results

### Fragments with similar pseudo-torsion angles can have dissimilar structures

In previous studies, pseudo-torsion angles (θ and η) were employed to classify all fragment nucleotide conformers ^19, 23^. If tri-nucleotide conformations are classified according 5° bins for η and θ, one would see a nearly complete coverage of all possible grid points as shown in Supplementary Fig. 1 (particularly for C3’endo conformation). However, two fragments with nearly identical pseudo-torsion angles do not necessarily have the same structure. Two examples are illustrated in Fig. 2. In Fig. 2a, two tri-nucleotide structures (derived from the nucleotides 17-19 of a riboswitch (2kxm_A) and 2581-2583 of a ribosome (7pwo_1), respectively) have nearly the same pseudo-torsion angles (η angles are 86.4° versus 88.1° and θ angles are -145.7° versus -149°).

**Figure 2.**
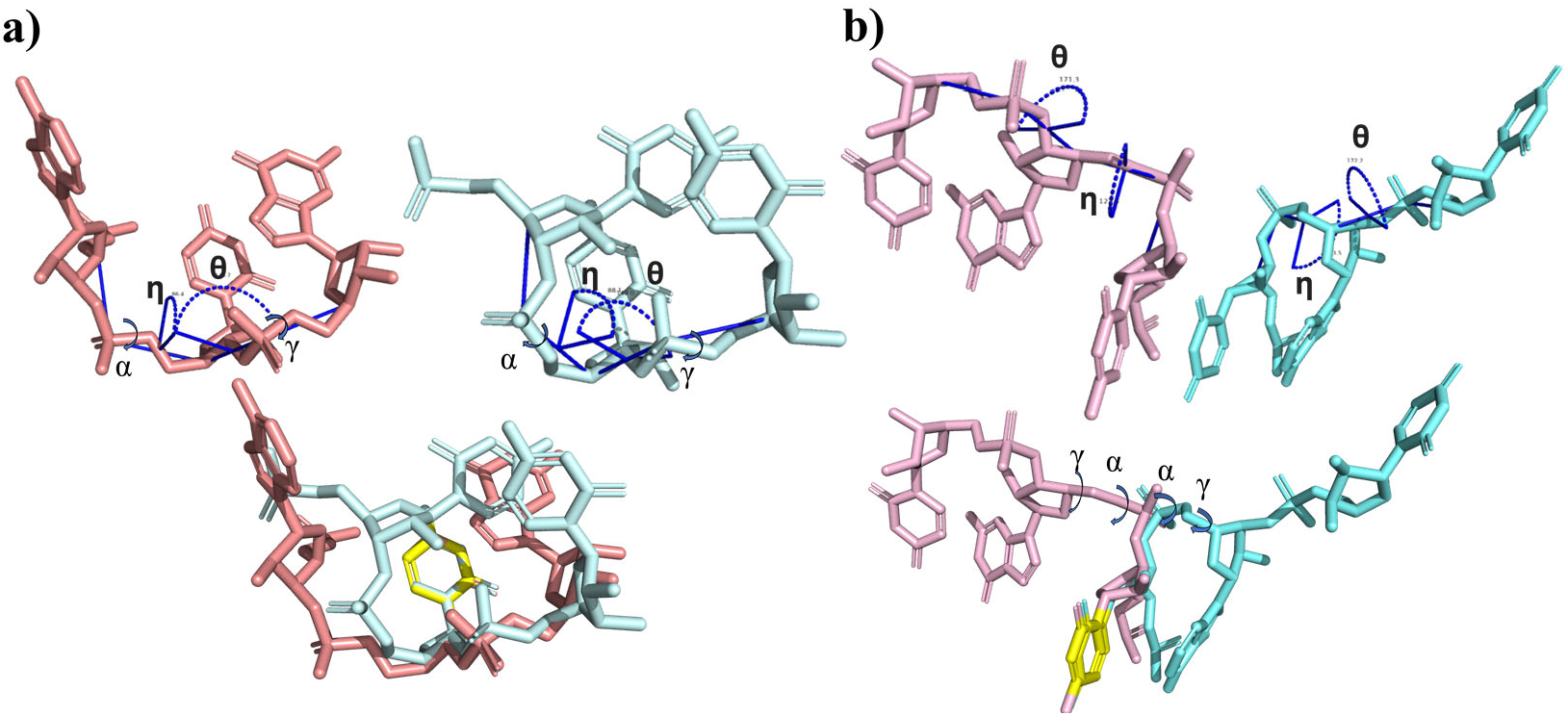
Two examples that tri-nucleotide conformers share the similar pseudo torsion angles but having large structural differences. The structural alignment shown was based on aromatic ring colored with yellow. a) Structural comparison between the fragment of nucleotides 17-19 of a riboswitch (2kxm_A, C3-endo sugar pucker, in pink) and the fragment of nucleotides 2581-2583 of a ribosome (7pwo_1, C3-endo sugar pucker, in blue)). b) Structural comparison between the fragment of nucleotides 42-44 of a synthetic RNA (7pdu_B for C3’-endo, in pink) and the fragment of nucleotides 3150-3152 of a ribosome (8euy_1 for C2’-endo, in blue).

However, the RMSD values between these two fragments are 3.55Å for the B0S3 base-atom representation (Fig. 1g) and 2.44Å for the B6S0 backbone-atom representation (Fig. 1d). These two fragments would be considered as different structural fragments based on our 1 Å threshold (see Methods). One reason for this large difference is that the differences between the backbone gamma torsion angles about the C5_2_’-C4_2_’ bond (59.6° versus -29.5°) and the alpha torsion angles about the P_3_-O5_3_’ bond (-80.9° versus 84.0°) of the two conformers are nearly 90° and 160°, respectively. Similarly, in Fig 2b, the structures of nucleotides 42-44 of a synthetic RNA (7pdu_B for C3’-endo) and nucleotides 3150-3152 of a ribosome (8euy_1 for C2’-endo) share nearly identical pseudo-torsion angles (122.5° versus 123.5° for η and 171.3° versus 172.2 for θ).

However, the B0S3 base frame RMSD and B6S0 backbone frame RMSD values between the two fragments are 3.13Å and 2.96Å, respectively. This is in part caused by the fact that the differences between alpha angles about the P_2_-O5_2_’ and the gamma angles about the C5_2_’-C4_2_’ of the two conformers are near 170° (174.3° versus -20.7° for alpha and -176.7° versus 13.6° for gamma). Thus, using two backbone pseudo torsion angles alone is insufficient to describe the backbone structural space of RNA fragments.

### Comparison between reduced representations

We measure whether a reduced representation is a good approximation to the full-atom representation by examining the correlation between RMSD values from the reduced representation and those from the full atom representation. We employed 8755 representative three-nucleotide fragments according to the mixed backbone-sidechain B2S1 representation (Fig. 1e) and obtained the RMSDs values for all studied representations. Fig. 3 shows the results for 100, 000 points randomly selected from all pairwise alignments. The highest correlation coefficient is between the RMSD value of the B2S1 frame and that of the full-atom representation (pcc=0.969). Other correlation coefficients range from 0.582 between all-atom and B0S3 representations, 0.659 between all-atom and B6S0 representations, and to 0.767 between all-atom and B1S2 representations. Thus, while base pairing drives RNA folding, it requires more than base atoms for characterizing the whole structural space of RNA.

**Figure 3.**
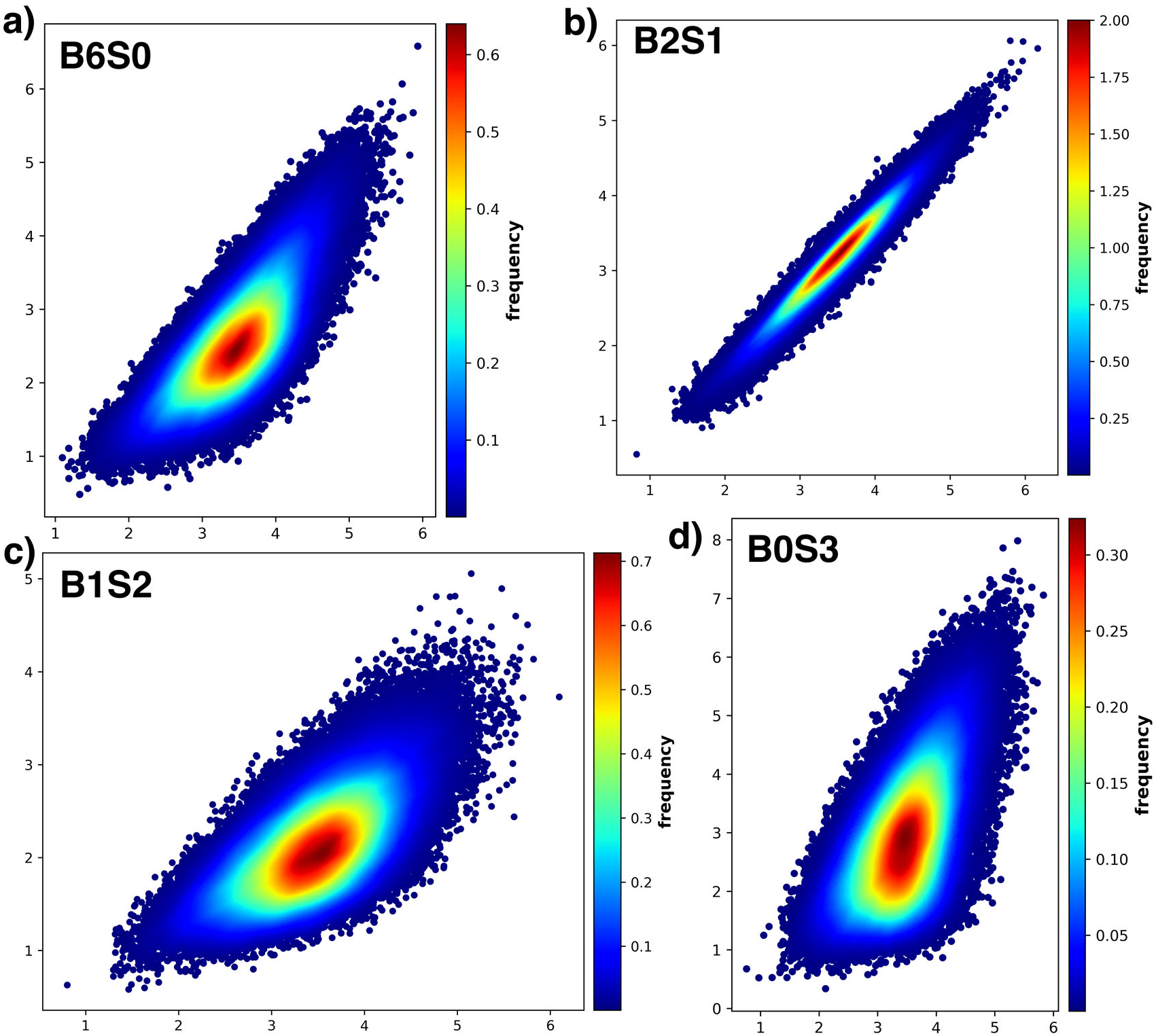
The correlation between all-atom RMSDs and those from other representations. a) the backbone frame (B6S0). b) the backbone-base-mixed frame (B2S1). c) the backbone-base-mixed frame (B1S2). d) the base frame (B0S3). The vertical ordinates and horizontal ordinates indicate full atom RMSD and frame RMSD, respectively.

### Clustering of protein and RNA trimeric fragments

Fig. 4a shows the structural space of RNA fragments according to the cumulative number of representative trimeric fragments by using protein fragments as a control. The number of increased protein fragments has been less than 1% since year 2015 (Fig. 4b). (The year 2015 is much later than what was reported^3^ because we employed a non-redudant set of proteins at 30% sequence identity cutoff for reducing the computational time.) By comparison, only the full-backbone frame (B6S0) achieved this 1% threshold at year 2023. However, the reach of this threshold for B6S0 is unlikely an indication of completeness, as it is a single datapoint and the increase in year 2022 is as large as 8.7%. Moreover, all other indicators have significant increase in new structural fragments in year 2023, particularly for the three-atom representation B2S1. The increase of fragments in the B2S1 representation is as large as 6% in 2023.

**Figure 4.**
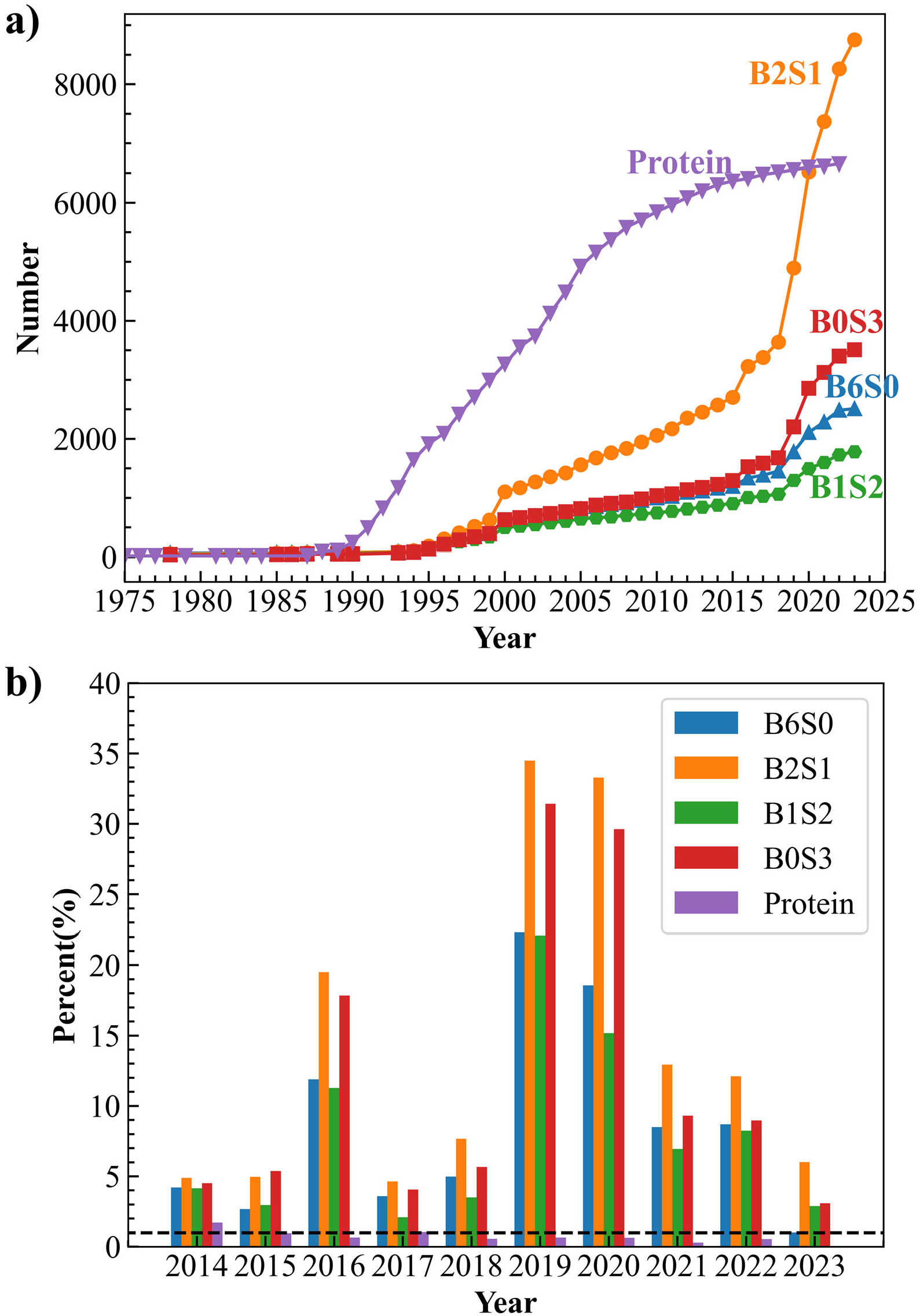
The cumulative number of structural fragments for proteins and RNA fragments as a function of the year (a) and the fraction of newly added structural fragments for every year compared to previous years.

We observed that there were sudden growths in the number of RNA fragments in years 2000, 2019, and 2020. The first high-resolution complete atomic structure of ribosomal subunit (pdb id: 1ffk) contributes to the year 2000 jump ^24^. Two new ribosome structures (70S ribosome complex with erythromycin (6s0x) ^25^ and 50S ribosome bound to compound 40e (6pcs) ^26^) are responsible most to the number of new structural fragments in 2019. Another ribosome structure (yeast 80S ribosome complex, 6woo) in 2020 leads to a large increase in new fragment structures. Thus, the completeness of RNA structure space might be disrupted if a new previously unseen RNA structure is found.

We also clustered all di-, tetra-, and penta-nucleotide conformers based on B2S1 and B0S3 frames. The trend of the number of these fragments are shown in Fig 5. The number of 4-mer and 5-mer fragments increases exponentially over the time. For 2-mer, it appears to reach the completeness for B0S3 since 2021, but not for B2S1 (an increase of new structure fragments by 2.4% in 2023).

**Figure 5.**
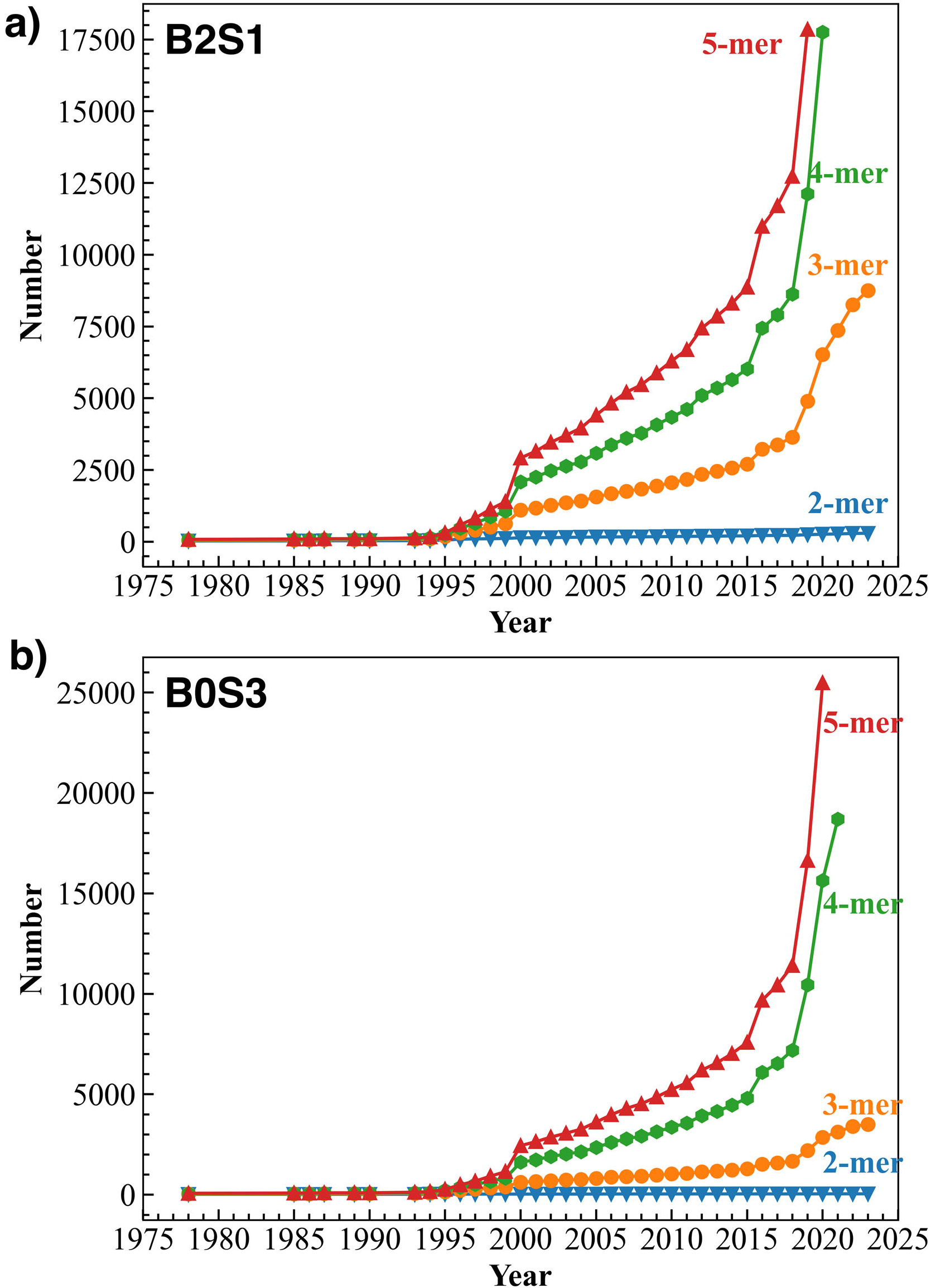
The cumulative number of structural fragments at the level of two, three, four, and five consecutive nucleotides as a function of the year. (a) the B2S1 representation; (b) the B0S3 representation.

We further analyzed all representative conformers based on their secondary structures. As is shown in Fig 6, the trimeric fragments located in loop (L) and mixed helix-loop (HL) regions increases over the time for both B2S1 and B0S3 representations. The increase in 2023 is 4.3% for HL and 7.0% for Loop in the B2S1 representation and 3.1% for HL and 4.0% for Loop for the B0S3 representation. By comparison, the trimeric fragments in helix regions seem to reach completeness for the B2S1 representation since the year 2022 and for the B0S3 representation since the year 2016, consistent with the simpler structural patterns in the helical region.

**Figure 6.**
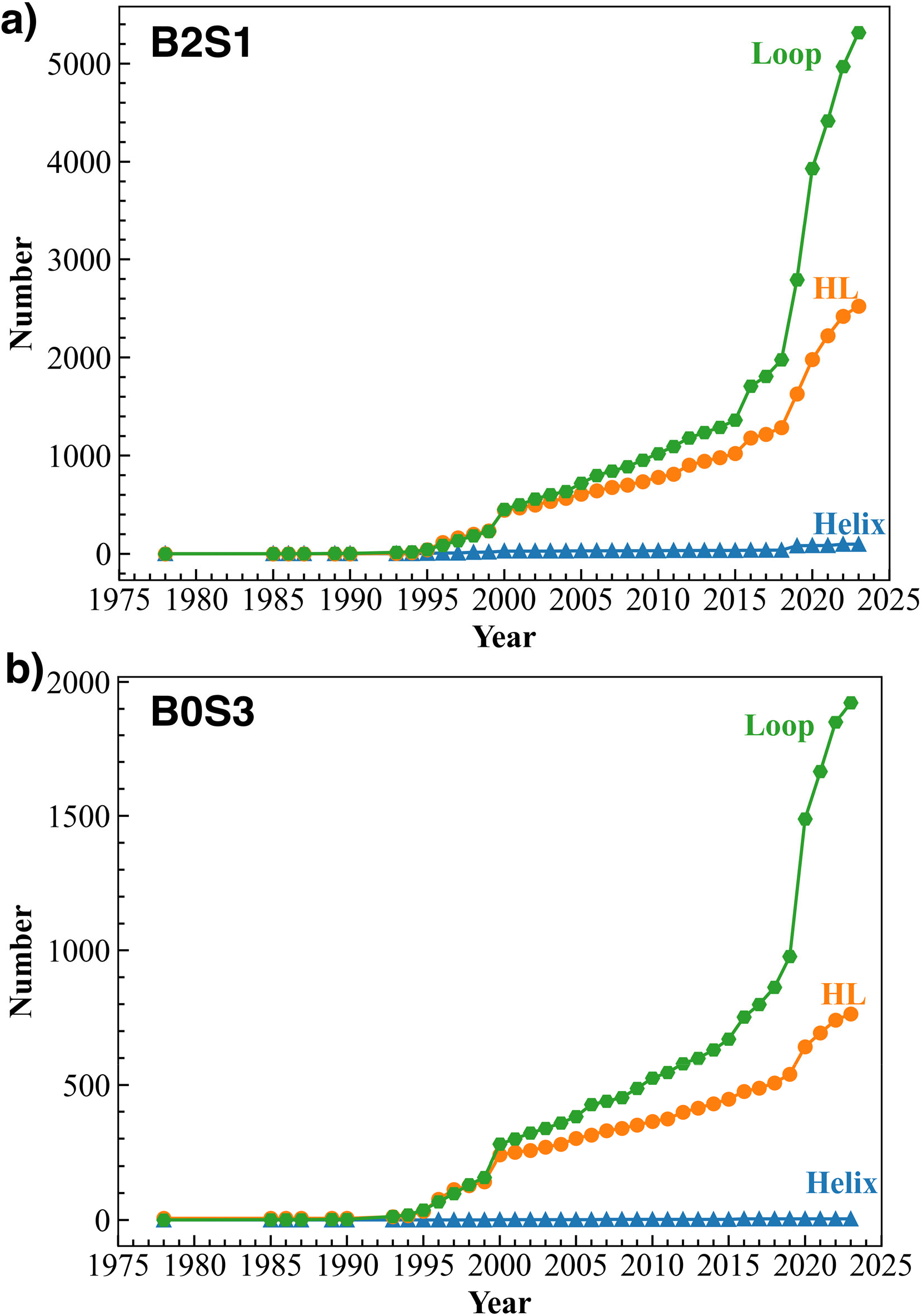
The cumulative number of tri-nucleotide structural fragments located on helix, loop, and helix-loop regions (HL) as a function of the year. (a) the B2S1 representation; (b) the B0S3 representation.

### Classification based on the trinucleotide sequence

Given the highly similar physio-chemical properties of bases, one would expect that all sequences had a similar number of fragments once the structural space of the fragments reached completeness. As shown in Fig S3, the number of fragment structures for the same trinucleotide sequence is uneven, ranging from 300 to 800, another indication of incompleteness in fragment structures. Even for the two sequences with the most structural fragments (GGA and GCC), the completeness is not yet reached, as shown in Fig S4.

## Discussion

To understand why predicting RNA structures remains a highly challenging problem, we examined the fragment structures of RNAs at dimeric, trimeric, tetrameric, and pentameric levels. Clustering these structures according to the RMSD values, we found that more than 1% fractions of new structural fragments continue to be found in newly solved RNA structures even at the di-nucleotide level. Although a slowdown in the increase of new fragments was observed for dimeric and trimeric fragments even witgh <1% increase for some reduced representations, an exponential increase is observed in new tetrameric and pentameric fragments. Thus, the structural space of RNAs observed is far from complete.

The incompleteness in the space of fragment structures is further supported by the observation that the number of trimeric structure fragments for RNAs (2510 according to the B6S0 backbone representation) is far smaller than that found for proteins (6652, Fig. 4). This result is opposite to our expectation because the RNA backbone has more degree of freedom (6 torsion angles) than the protein backbone (3 torsion angles with one nearly fixed at 0 or 180 degree). Moreover, the uneven number of observed structural fragments for different three-nucleotide sequences is consistent with the incompleteness of the structural space observed, as similar physio-chemical properties of nucleobases should have nearly the same number of structural fragments, regardless of sequence combinations.

One might argue that there is a slowdown in the growth of new fragment structures at least at the dimer and trimer level. This could be temporary. New structural space can be expanded rapidly if a new structure was solved, as it happened around years 2000, 2019, and 2020 (Fig. 4). Thus, we might observe another significant expansion if the structure for a new class of RNA molecules is solved in the future.

How to increase the chance of finding more new structure fragments? We examined the relation between new fragments found and the number of RNA structures in non-redundant sequences deposited in the PDB each year. As Fig. S4 shows, there is a moderate positive correlation (range from 0.5 to 0.61) between the number of new structural fragments and the increased number of deposited structures in non-redundant sequences. Thus, solving more structures is an effective way to expand the structural space.

To locate the best reduced representation for RNA structures, we examined four different frames (backbone only, sidechain only and two combinations of sidechains and backbone atoms). Different representations were employed previously for RNA structure prediction by deep learning. The B2S1 representation was employed in DRfold ^9^, whereas the B1S2 representation was employed in E2Efold-3D ^8^. Our result indicates that the B2S1 representation can better characterize the overall structure of an RNA, given its RMSD values having near perfect correlation to the full-atom RMSD (Fig. 3). However, the B2S1 representation has the highest number of structural fragments whereas B1S2 has the lowest number of structural fragments, which is even much lower than the number from the pure backbone (B6S0) or the pure base representation (B0S3). This suggests that the B1S2 frame has the least structural variation (or, structurally most stable). Thus, the B1S2 frame may be the best for model building for deep learning algorithms or fragment-based assembly.

## Methods

### Structure datasets

We downloaded all structures containing proteins or RNAs from the PDB database ^2^ as of Mar 5^th^, 2024. We extracted the structures solved by NMR structures, X-ray diffraction, and cryo-electron microscopy with resolution better than 3 Å. All structures released before Jan 1^st^, 2024 were employed for analysis. There are 1089 RNA chains and 31926 protein chains after removing redundance by CD-HIT ^27^ with sequence identity cut off at 0.8 and 0.3, respectively. Time stamps of these structures were recorded so that we can construct time-dependent fragment libraries.

### Clustering of structural fragments

The most employed torsion angles for characterizing backbones are ϕ and ψ (Fig 1(a)) for proteins ^28^ and θ and η (Fig 1(c)) for RNAs ^19^. For RNA, θ and η are rotational angles around the pseudo-bonds between P_i_ and C4’_i_ and between C4’_i_ and P_i+1_, respectively. They were employed for a coarse-grained representation of a RNA backbone.

Structural similarity between two fragments can be measured according to root-mean-squared distance (RMSD) between the frame atoms. For proteins, three backbone atoms are often employed as the frame atoms (N, Cα, C, Fig. 1(c)). RNA structures are made of the sugar-phosphate backbone and nitrogenous base sidechains. We have examined several representations of fragment structures: all backbone atoms (P, O5’, C5’, C4’, C3’, and O3’, denoted as B6S0, Fig. 1(d)), key sidechain atoms (N9, C8, C4 for A and G, N1, C6, C2 for C and U, denoted as B0S3, Fig. 1(g)), a combination of two backbone and one sidechain atoms (C4’, P and N1 for C/U or C4’, P and N9 for A/G, denoted as B2S1, Fig. 1(e)), and a combination of one backbone and two sidechain atoms (C4’, C1’ and N1 for C/U or C4’, C1’ and N9 for A/G, denoted as B2S1, Fig. 1(f)). The RMSD values of these reduced representations are compared to the full-atom RMSD calculated based on all heavy atoms (P, OP1, OP2, O5’, C5’, C4’, O4’, C3’, O3’, C2’, O2’, C1’, N9/N1, C8/C6, C4/C2).

We followed a previous study for fragment structure clustering ^3^ by calculating RMSD values between the frame atoms of the two fragment structures. First, we clustered all fragments with a threshold of 0.5Å and obtained the representative fragments for each structure cluster and the number of fragment structures for each structure cluster. Second, all representative fragments of all structures in each year are further clustered with a threshold of 0.5Å and obtained the representative fragments and the number of fragment structures for each cluster. Afterward, we compared the representative fragments in each year to all representative fragments in previous years and counted the number of the fragments with RMSD > 1 Å as the new structural fragments found in each year. The Kabsch algorithm ^29^ was utilized to calculate all pairwise RMSD values and we applied hierarchical clustering implement by python to cluster all fragments based on their RMSD values. Same as the previous study ^18^, the threshold for structure clustering is 1Å for both protein and RNA fragments and the completeness of the structural space is defined as reached if the percentage of newly added fragments is less than 1% of the representative fragments from all previous years^3^.

## Competing interests

The authors have declared no competing interests.

## Acknowledgments

We thank Mr. Qianzhi Sang for assistance in plotting Figure 2. This work is supported by Natural Science Foundation of China [22350710182]. The authors gratefully acknowledge that the High Performance Computing Cluster at Shenzhen Bay Laboratory was involved in completing this study.

## Author contributions

**X**.**H**.: Data curation, Writing original draft, Writing - review & editing, Visualization. **J**.**Z**.: Conceptualization, Writing - review & editing, Supervision. **Y**.**Z**.: Conceptualization, Resources, Writing - original draft, Writing - review & editing, Supervision, Project administration, Funding acquisition. All authors have read and approved the final manuscript.

## Supplementary Materials

**Figure S1.**
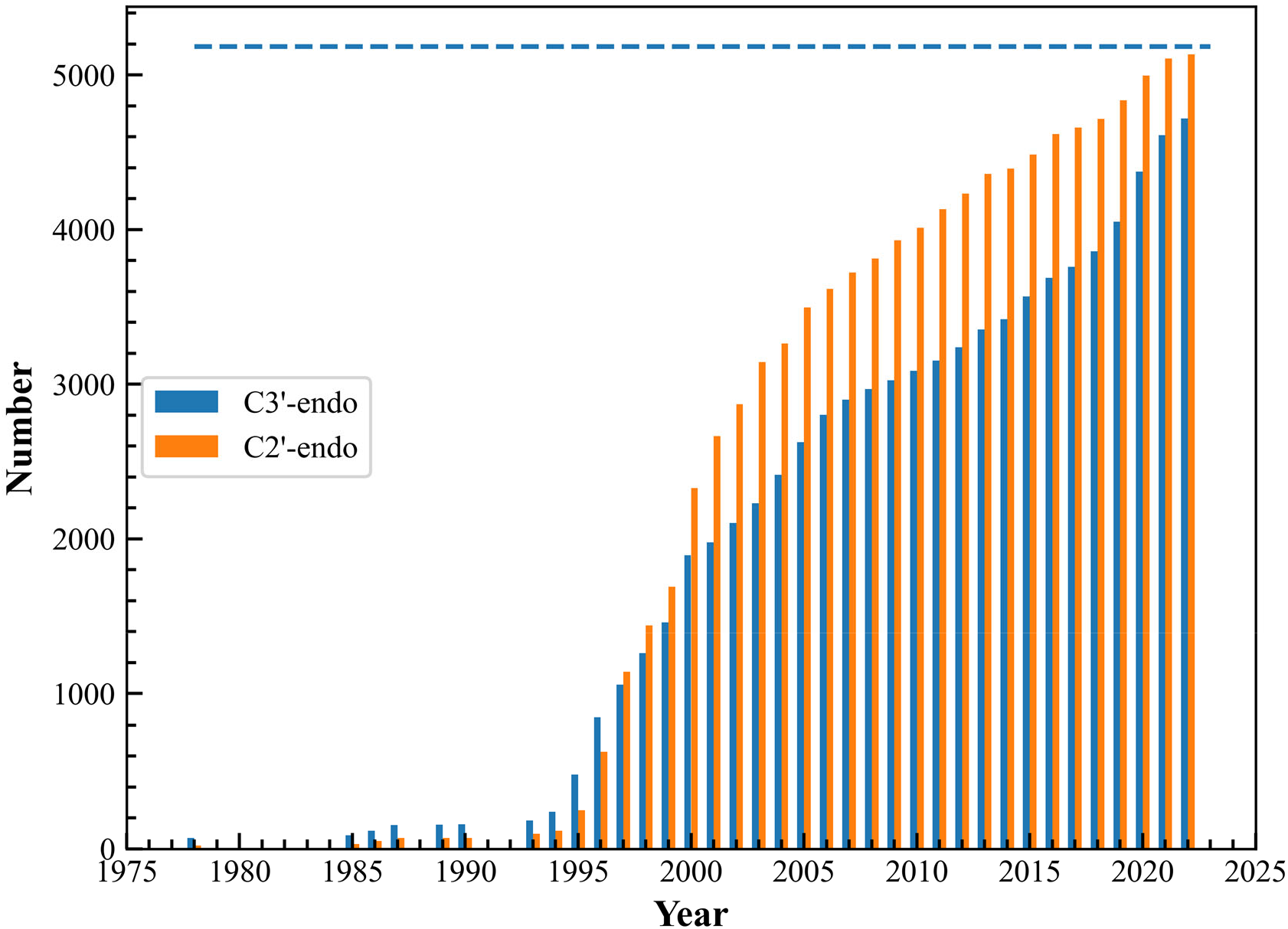
The total number of clusters as a function of the year based on a 5° bin in two pseudo torsion angles. The maximum total is 5184 (72×72) bins.

**Figure S2.**
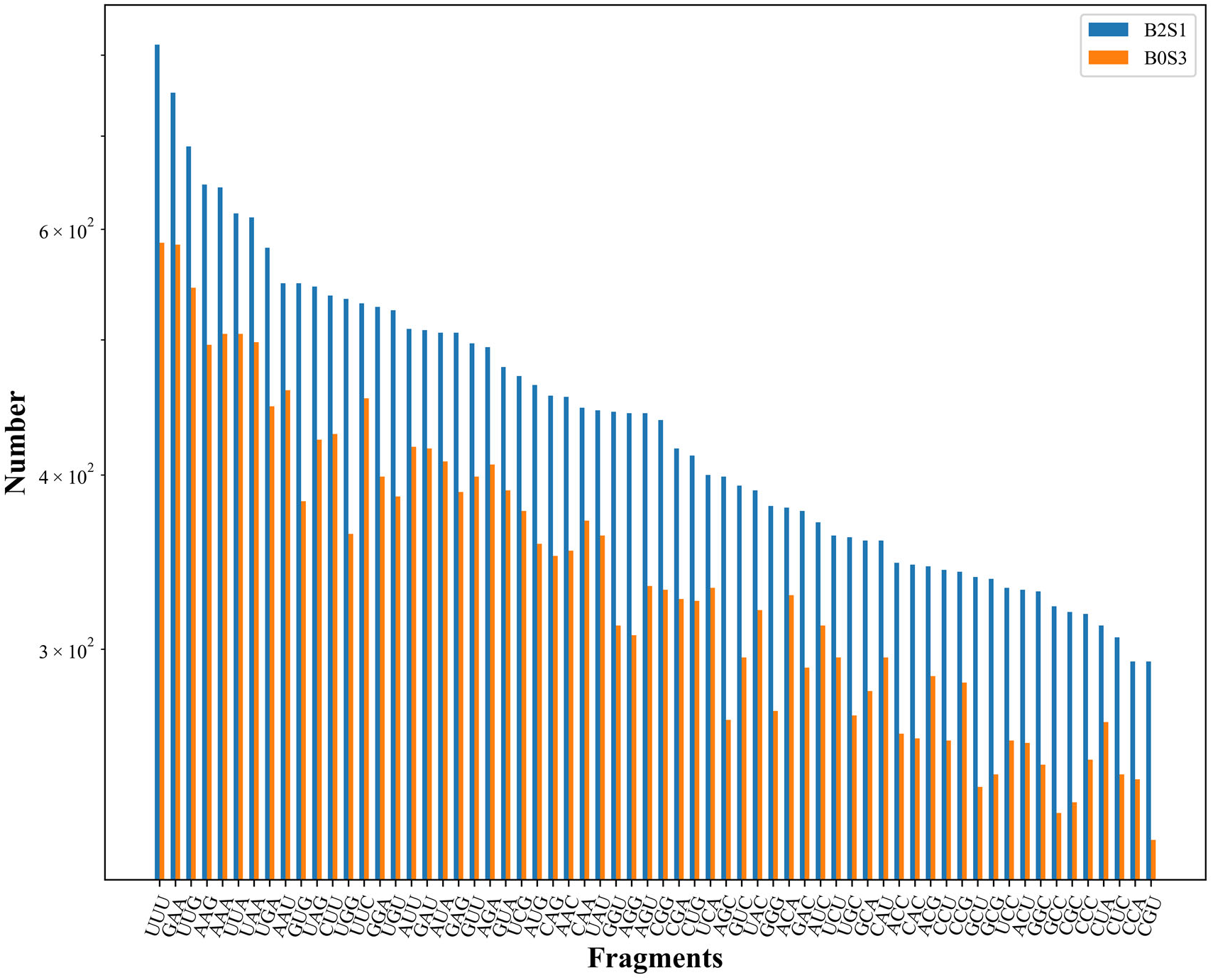
The number of structural fragments based on different trinucleotide sequences. All structural fragments were clustered based on either B2S1 or B0S3 representations as labeled.

**Figure S3.**
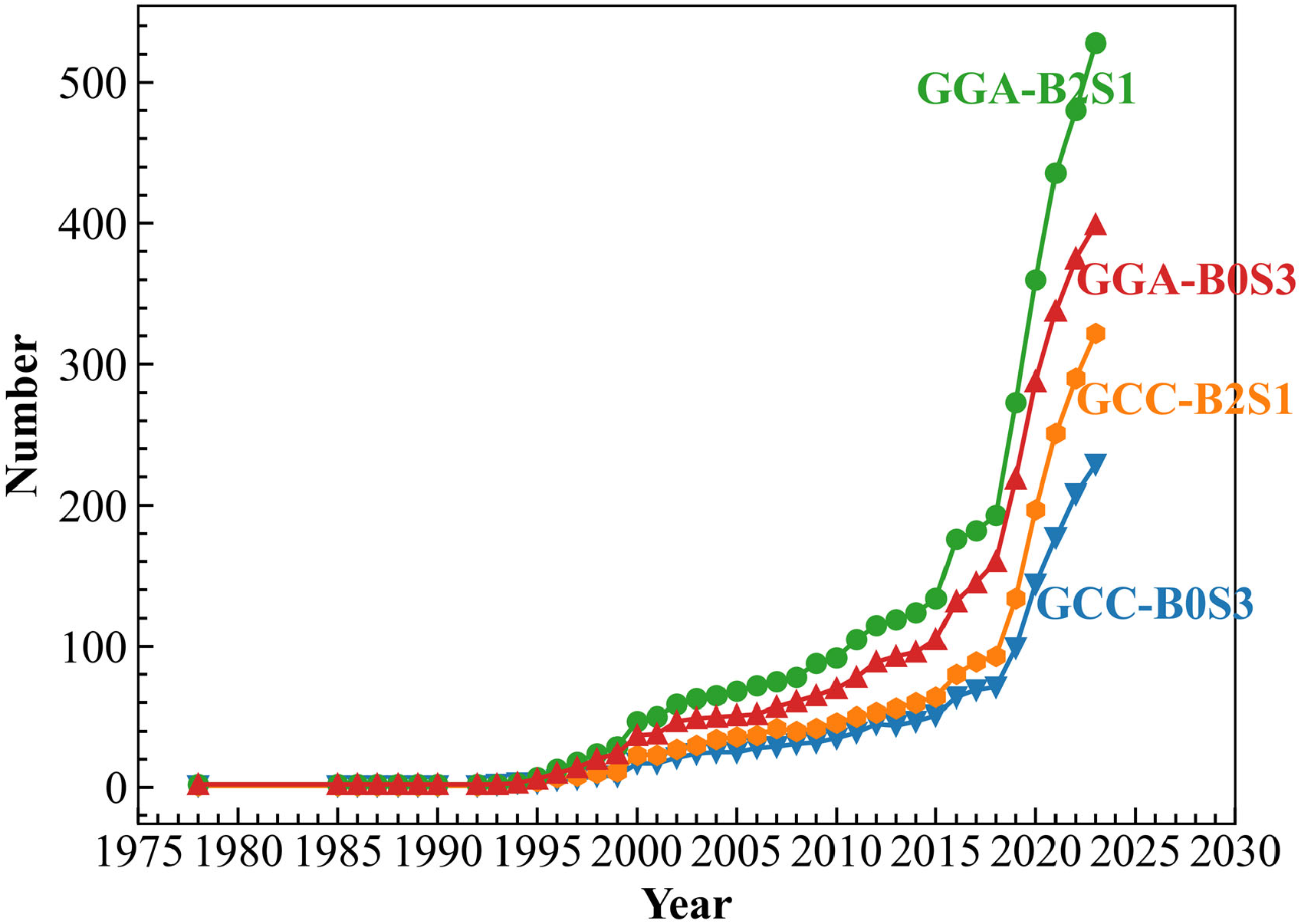
The number of structural fragments for two trinucleotide sequences (GCC and GGA) clustered by the B2S1 and B0S3 representations as a function of the year.

**Figure S4.**
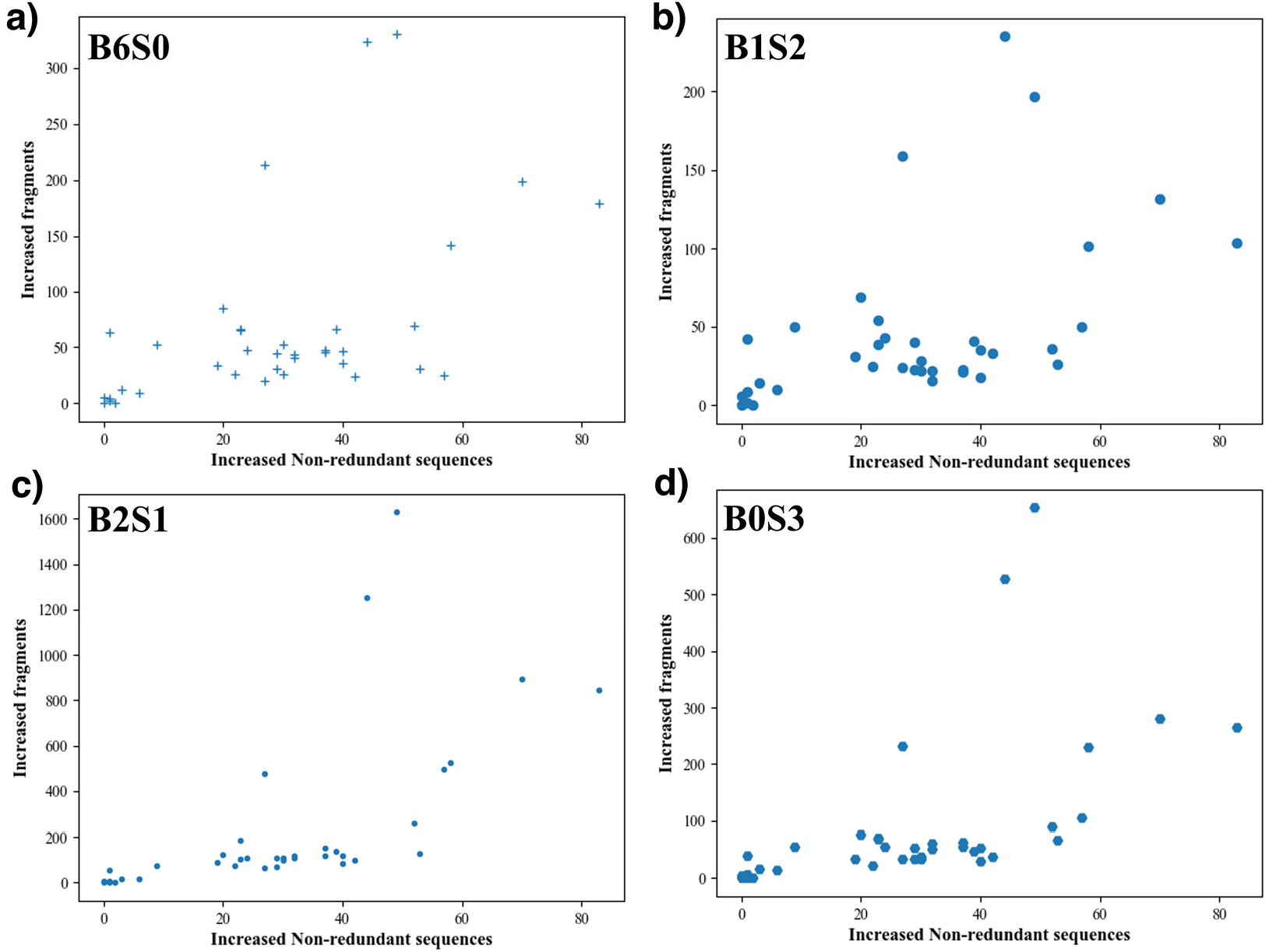
The number of newly added structural fragments at a given year versus the number of deposited RNA structures in non-redundant sequences for the same year. a) B6S0 representation (Pearson R: 0.53, P value:8.1e-4). b) B1S2 representation (Pearson R: 0.50, P value:1.7e-4). c) B2S1 representation (Pearson R: 0.61, P value:5.5e-5). d) B0S3 representation (Pearson R: 0.53, P value: 6.7e-4).

